# Time-resolved secretome analysis of three *Colletotrichum* species identifies copper radical alcohol oxidases for the production of fatty aldehydes

**DOI:** 10.1101/2021.06.09.447826

**Authors:** David Ribeaucourt, Safwan Saker, David Navarro, Bastien Bissaro, Elodie Drula, Lydie Oliveira Correia, Mireille Haon, Sacha Grisel, Nicolas Lapalu, Bernard Henrissat, Richard J. O’Connell, Fanny Lambert, Mickaël Lafond, Jean-Guy Berrin

## Abstract

Copper Radical Alcohol Oxidases (CRO-AlcOx), which have been recently discovered among fungal phytopathogens are attractive for the production of fragrant fatty aldehydes. To investigate the secretion of CRO-AlcOx by natural fungal strains, we undertook time-course analyses of the secretomes of three *Colletotrichum* species (*C. graminicola, C. tabacum* and *C. destructivum*) using proteomics. The addition of a copper-manganese-ethanol mixture to *Colletotrichum* cultures unexpectedly induced the secretion of up to 400 proteins, 29-52% of which were carbohydrate-active enzymes (CAZymes), including a wide diversity of copper-containing oxidoreductases from the auxiliary activities (AA) class (AA1, AA3, AA5, AA7, AA9, AA11-AA13, AA16). Under these specific conditions, while a CRO-glyoxal oxidase from the AA5_1 subfamily was among the most abundantly secreted proteins, the targeted AA5_2 CRO-AlcOx were secreted at lower levels, suggesting heterologous expression as a more promising strategy for CRO-AlcOx production and utilization. *C. tabacum* and *C. destructivum* CRO-AlcOx were expressed in *Pichia pastoris* and their preference toward both aromatic and aliphatic primary alcohols was assessed. The CRO-AlcOx from *C. destructivum* was further investigated in applied settings, revealing a full conversion of C6 and C8 alcohols into their corresponding fragrant aldehydes.

**IMPORTANCE:** In the context of the industrial shift toward greener processes, the biocatalytic production of aldehydes is of utmost interest owing to their importance as intermediates in preparative chemistry and for their use as flavors and fragrances ingredients. In the search for new biocatalysts, CRO-AlcOx have the potential to become platform enzymes for the oxidation of alcohols to aldehydes. The use of crude fungal secretomes is often seen has an appealing approach by industries since alleviating various costs pertaining to biocatalysts production. However, the secretion of CRO-AlcOx by natural fungal strains has never been explored. This study showed that *Colletotrichum* species can secrete a broad diversity of copper-containing enzymes, but only little amount of CRO-AlcOx. Thus, recombinant expression remains the most promising approach. The potential of CRO-AlcOx as biocatalyst for flavor and fragrance applications was confirmed through the production of two new enzymes with activity on fatty alcohols.

## INTRODUCTION

The development of a new technological paradigm to support the shift from a fossil-based economy to a greener and more circular economy is at the heart of today’s bioeconomic challenges (1). Biotechnology is one of the main levers in this ongoing industrial transformation (2). The development of biotechnology largely relies on biocatalysis and thus require the discovery, control and engineering of a variety of enzymes and microorganisms (3). The fungal kingdom represents a vast and untapped reservoir of biocatalysts (4) but to date, the enzymatic potential of filamentous fungi has mostly been studied in the frame of plant cell wall degradation (5–7). Fungal secretomics is a powerful approach to study the wide diversity of enzymes secreted and to formulate enzymatic cocktails with biotechnological relevance (8–10). Using suitable inducers, some fungal strains can secrete high amounts of oxidoreductases (11) including copper-containing enzymes. For instance, hypersecretory fungal strains from the *Pycnoporus* genus have been selected to produce laccases and use them in applied settings (12, 13).

Most of the copper-dependent enzymes are classified within the Auxiliary Activities (AA) class (14) in the CAZy database (www.cazy.org) (15). The founding member of copper radical oxidases (CROs), a galactose oxidase (EC 1.1.3.9), was discovered in 1959 from the secretome of *Fusarium graminearum* (16, 17). In 2013, fungal CROs have been classified within the AA5 family and further classified into subfamilies AA5_1 and AA5_2. AA5_1 contains glyoxal oxidases (GLOX - EC 1.2.3.15) that catalyze the oxidation of aldehydes to carboxylic acids, and AA5_2 gathers enzymes that oxidize the alcohol function of diverse substrates (e.g., galactose, galactose-containing polysaccharides, aromatic and aliphatic primary alcohols) to generate the corresponding aldehydes. More recently, the AA5_2 subfamily was shown to be more diverse than expected, encompassing notably various alcohol oxidases (CRO-AlcOx; EC 1.1.3.13) originating from the phytopathogen ascomycete fungi *Colletotrichum* and *Magnaporthe* spp. (18–20). CRO-AlcOx are monocopper, and otherwise organic co-factor free enzymes that catalyze the oxidation of alcohols into aldehydes (with the concomitant reduction of O_2_ to H_2_O_2_). They offer appealing perspectives to produce aldehydes (21), which are indispensable intermediates in many chemical pathways (22, 23) and valuable ingredients for the flavor and fragrance industries (24, 25). In particular, saturated long-chain aldehydes (fatty aldehydes) are widely used for their citrus scent (26, 27), but are currently derived from fossil-based chemistry (28, 29) or extracted from natural plant materials, the supply of which is threatened by emerging diseases (30, 31). Therefore, alternative green and/or natural production pathways must be investigated (32).

To date, only pure recombinant forms of CRO-AlcOx have been harnessed in biocatalytic applications (18, 19, 21, 33) and no information on their secretion by natural fungal strains is available. Determining whether CRO-AlcOx encoding fungal strains can secrete these enzymes under laboratory conditions is of strong interest for enzyme selection and technological applications. Noteworthy, aside from the scope of pathogenesis (34–36), very few studies investigated the enzymatic biotechnological potential of *Colletotrichum* spp. (37–40). The use of natural fungal strains could be advantageous to identify and then use potential naturally secreted enzyme partners. In particular, catalase and peroxidase are both required to sustain *in vitro* the activity of isolated CRO-AlcOx and other AA5 (21, 33, 41, 42). Catalase acts as a H_2_O_2_ scavenger while peroxidase is required for CRO-AlcOx activation (20, 33).

In this context, we investigated the ability of several fungal strains from the *Colletotrichum* genus to secrete copper-containing enzymes with a focus on CRO-AlcOx. We report here an integrated study, ranging from fungal time-course secretomics to recombinant enzyme characterization, and biocatalytic application of CRO-AlcOx enzymes to produce fatty aldehydes for the flavor and fragrance industries.

## RESULTS

### Selection of fungal strains and time-course secretomic study

To investigate the secretion of copper-containing enzymes including AA5_2 CRO-AlcOx by natural fungal strains and their potential use for bioconversion of fatty alcohols to fragrant aldehydes, we selected three fungal phytopathogens from the *Colletotrichum* genus for which genomic information is available. The first obvious choice was *C. graminicola* because the three AA5_2 it encodes have all been previously biochemically characterized (18, 43, 44), one of them being the first, most studied, and to date, most active CRO-AlcOx, namely *Cgr*AlcOx (18, 21, 33). By performing a tblastn analysis of the *Cgr*AlcOx sequence against the genomes of some other *Colletotrichum* species we had access to, we identified two genes encoding AA5_2 in both *C. tabacum* and *C. destructivum*. For each strain, one gene encoded a putative AlcOx and the other one a putative aryl alcohol oxidase (AAO) (Figure S1). These enzymes will thereafter be named *Cta*AlcOx/*Cde*AlcOx and *Cta*AAO/*Cde*AAO for *C. tabacum* and *C. destructivum*, respectively. Based on the few studies reporting the detection in some Ascomycota fungal secretomes of a range of copper-containing enzymes from the AA class including the *Fgr*GalOx and other AA5 (45–47), we elaborated two liquid growth media: (i) “YG” (for “Yeast-extract, Glucose”), as a minimal medium, and (ii) ‘YG-CME” (for “YG-Copper, Ethanol, Manganese”), with the same composition as YG but supplemented with CuSO_4_, manganese and ethanol. A time-course analysis was performed on each strain in both media at three time points (day 3, day 7 and day 14) and proteomic analyses were performed on the 16 secretomes generated.

The quantification of total proteins revealed an increasing amount of proteins over time (except for *C. destructivum*) (Figure S2) while capillary electrophoresis analysis revealed distinct profiles over time and between fungal strains (Figure S3). By performing proteomic analyses of the secretomes using LC-MS/MS and mass matching, we detected a high number of secreted proteins (between 150 and over 400 different proteins, depending on the culture conditions; Supplementary Data Set; Figure 1A) encompassing a large diversity of enzymes. CAZymes accounted for ca. 29-52% of the total protein abundance in each secretome (Figure S4), with a notable high diversity of enzyme from the AA class (Figure 1B). Among them, a variety of laccases (AA1), peroxidases (AA2), flavo-oxidases (AA3) including cellobiose dehydrogenases (AA3_1-AA8), CROs (AA5), oligosaccharide oxidases (AA7), lytic polysaccharide monooxygenases (LPMOs; AA9, AA11, AA13, AA16) and pyrroloquinoline quinone-dependent oxidoreductase (AA12) were detected. To determine whether this high number and diversity of proteins detected was the results of cell lysis (7, 48), we evaluated the proportion of proteins predicted as not secreted – i.e., without signal peptide – and checked more specifically the abundance of some known intracellular proteins (i.e., DNA-polymerase, proteasome endopeptidase complex, citrate synthase, malate dehydrogenase, transketolase and ubiquitin-protein ligase) (Figure S5). Despite the significant number of proteins predicted without signal peptide, we did not observe any clear trend that would account for an accumulation of intracellular proteins in the secretomes over time. Notably, some proteins without signal peptide are known to be secreted through unconventional secretion pathways in fungi (48, 49).

**Figure 1:**
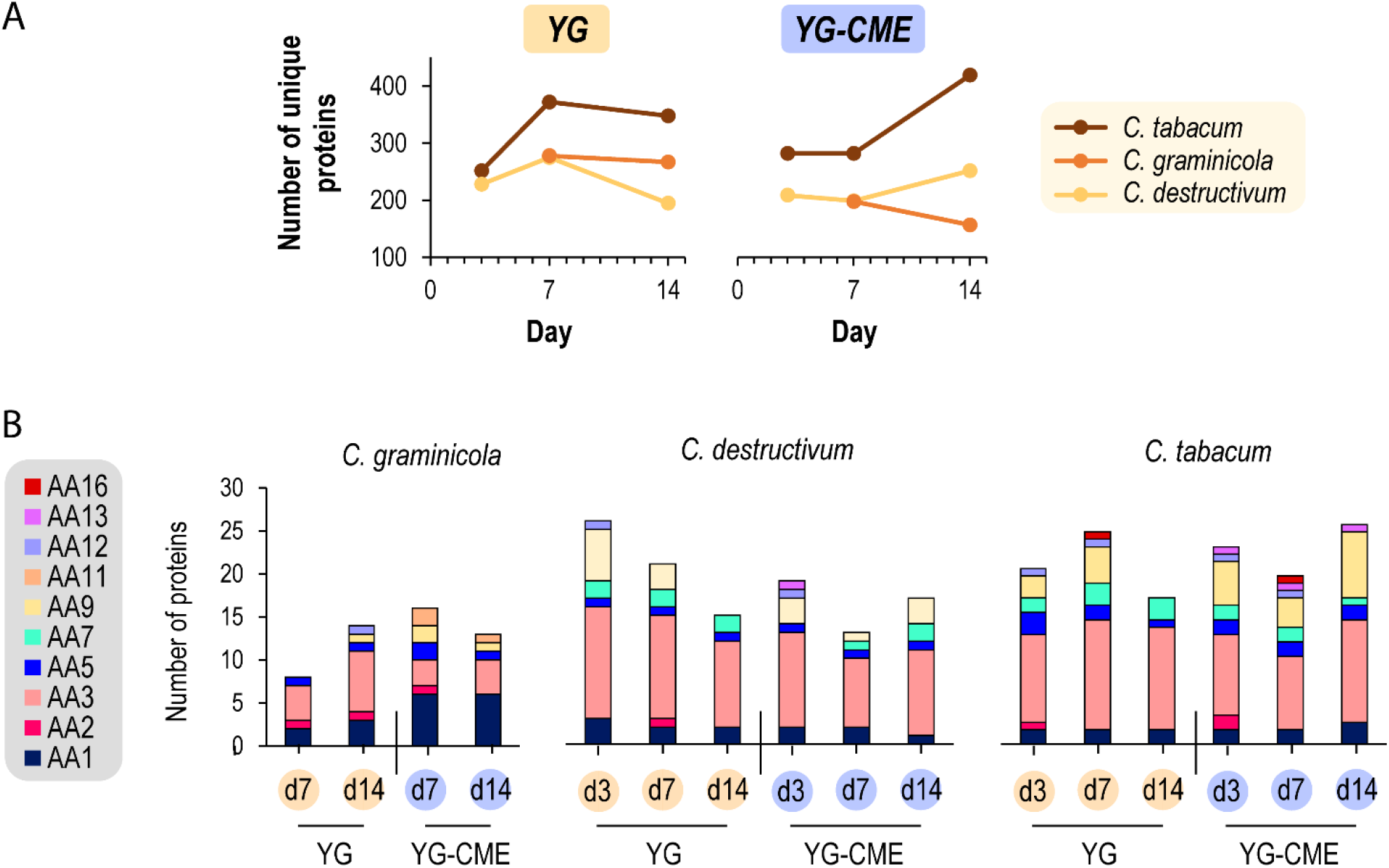
Overview of (A) the total number of proteins secreted over time in different media and (B) diversity of AA enzymes detected in the secretomes. Note: no samples were harvested at day 3 for *C. graminicola* due to insufficient growth. Abbreviations: CME, copper-manganese-ethanol; YG, minimal medium.

### Effect of Copper-Manganese-Ethanol supplementation on the secretome profiles

We then looked at the effect of supplementation with CME. At first glance, no clear effect was distinguishable on CAZymes diversity (Figures 1 & S4) but when we focused on copper-containing enzymes (Figure 2), a clear trend became apparent. Indeed, copper-containing enzymes were generally more abundant in the CME medium with a diversity of oxidases from the AA class including AA1 laccases, AA5_1 GLOX and AA9, AA11, AA13 and AA16 LPMOs. Furthermore, AA1_3, AA5_1 and AA9 were among the top-five most abundant enzymes in YG-CME for the *C. tabacum* and *C. destructivum* secretomes, while no AA enzymes were identified in the top-five ranking for the corresponding YG secretomes (Table S1). Remarkably, the presence of some AA9-CBM18 was detected in both the *C. tabacum* and *C. destructivum* secretomes (Figure 2). This association of a chitin-binding domain from the CBM18 family (mainly found associated to chitin-active enzymes) to an AA9 domain is intriguing since AA9 LPMOs have not yet been described to act on chitin substrates.

**Figure 2:**
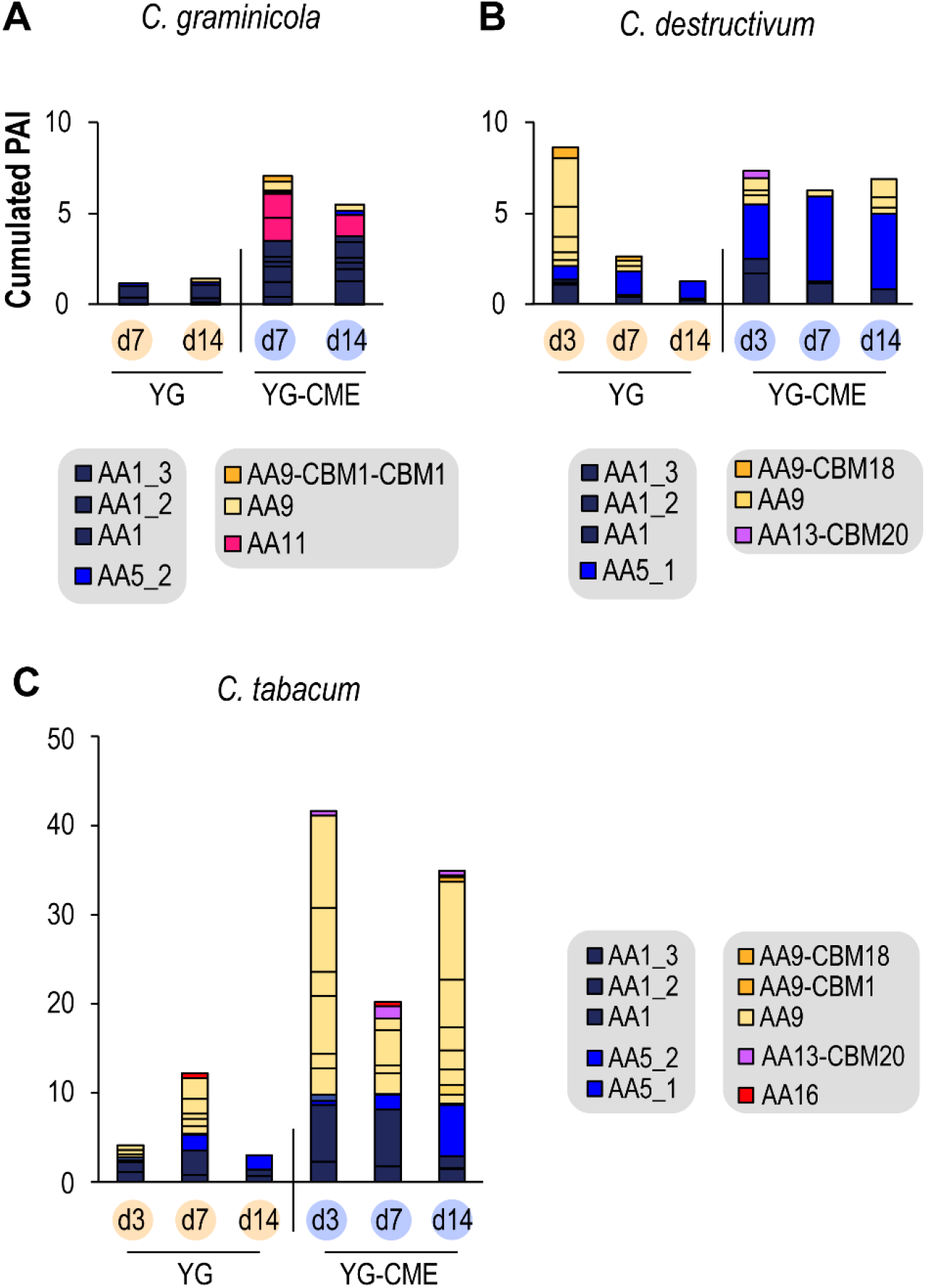
Abundance of copper-containing AAs in fungal secretomes from (A) *C. graminicola*, (B) *C. destructivum* and (C) *C. tabacum*. The Y-axis legend on panel A applies to all panels likewise (note the different scale for *C. tabacum*). For each enzyme family, the bar is subdivided according to the number of enzymes identified; for instance, *C. tabacum* at day 3 in YG-CME displayed six different AA9. PAI: Protein Abundance Index.

In addition, a diversity of Glycoside Hydrolases (GH), Polysaccharide Lyases (PL), Carbohydrate Esterase (CE) and AAs were detected exclusively in the CME medium (Figure 3). Among these enzymes, we noticed the intriguing presence of hydrolases (GH18, GH18-CBM18), oxidases (AA7, AA11) and esterases (CE4-CBM18) potentially active on chitin or chito-oligosaccharides. Remarkably, a multimodular protein with five consecutive CBM50 chitin-binding domains (“CBM50 (x5)” in Figure 3) but without any catalytic domain was detected in both the *C. tabacum* and *C. destructivum* secretomes. This protein is analogous to other chitin-binding proteins (i.e., LysM effector proteins) (see discussion).

**Figure 3.**
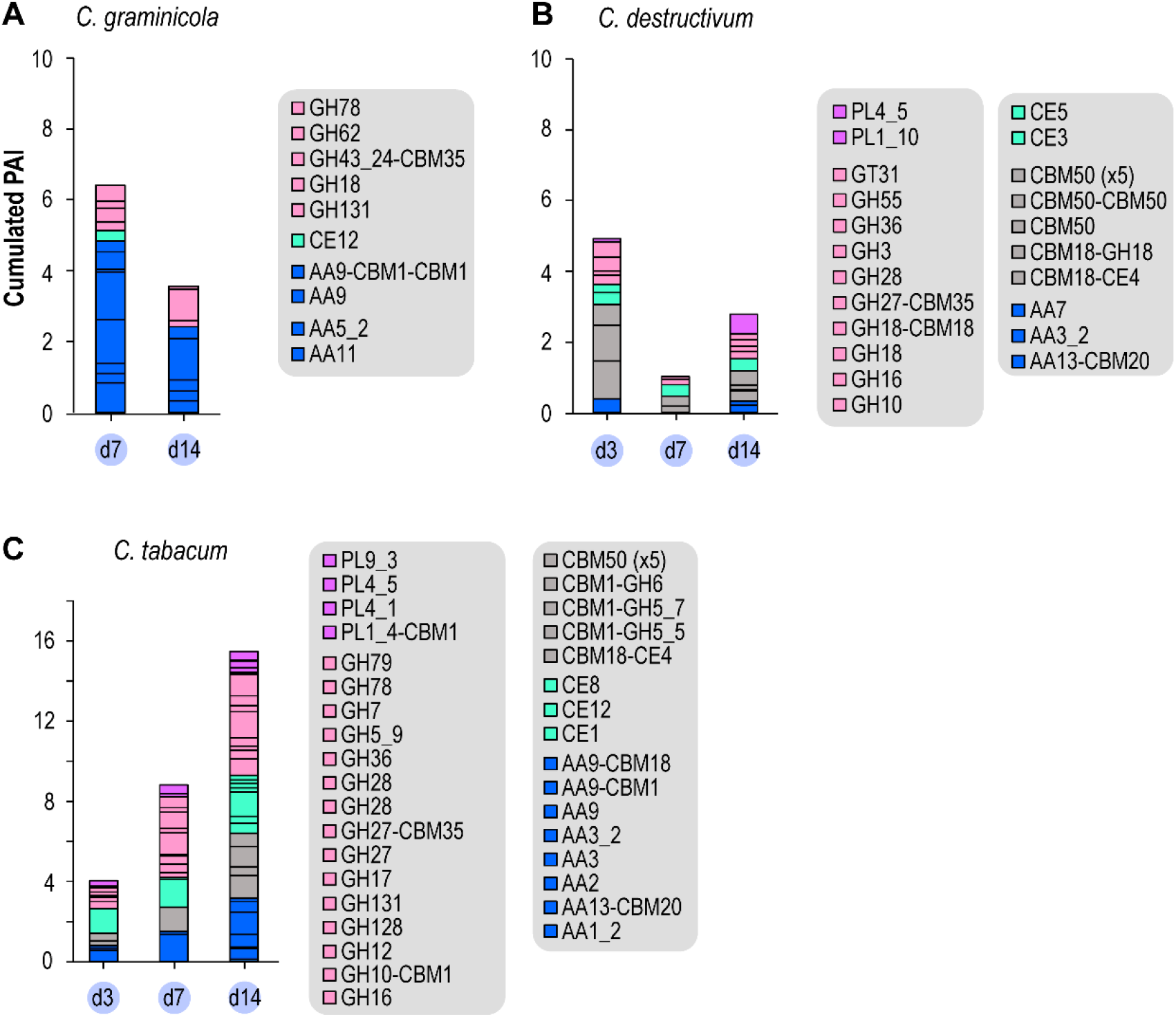
Abundance of CAZymes identified exclusively in the YG-CME media from (A) *C. graminicola*, (B) *C. destructivum* and (C) *C. tabacum*. The Y-axis legend on panel A applies to all panels likewise (note the different scale for *C. tabacum*). For each enzyme family, the bar is subdivided according to the number of enzymes identified. PAI: Protein Abundance Index.

### Presence of AA5 in the secretomes

We next focused our analysis on AA5 CROs. Interestingly, a putative GLOX from the AA5_1 subfamily was detected in *C. destructivum* secretomes (Figure 4), being the most abundant protein in the YG-CME secretome at day 7 (and the second most abundant at day 14; Table S1). The *C. tabacum* AA5_1 putative GLOX was also the third most abundant protein in the YG-CME secretome at day 14 (Table S1). We could also detect the occurrence of putative AlcOx and AAO from the AA5_2 subfamily in both *C. graminicola* and *C. tabacum* secretomes (Figure 4). However, their relative abundance was quite low (between 0.05-0.13% of the total Protein Abundance Index (PAI)) and they did not accumulate over time. No correlation with the presence of CME in the medium was seen except for the *C. graminicola* AAO. CROs are known to be activated by peroxidases (41, 50, 51). More specifically, the peroxidase partner responsible for the activation of CRO-AlcOx has been recently unveiled in fungal pathogens (20). Both genes occur as a tandem in the genomes of the *Colletotrichum* and *Magnaporthe* species harboring an AlcOx, and are co-expressed *in vivo* prior to the plant-penetration stage (20). While some peroxidases from the AA2 family (see Figure 2) were found in some of the secretomes, the specific peroxidase partner of the AlcOx was not detected reinforcing the authors claims that interaction with plant is needed for genes co-expression (20).

**Figure 4.**
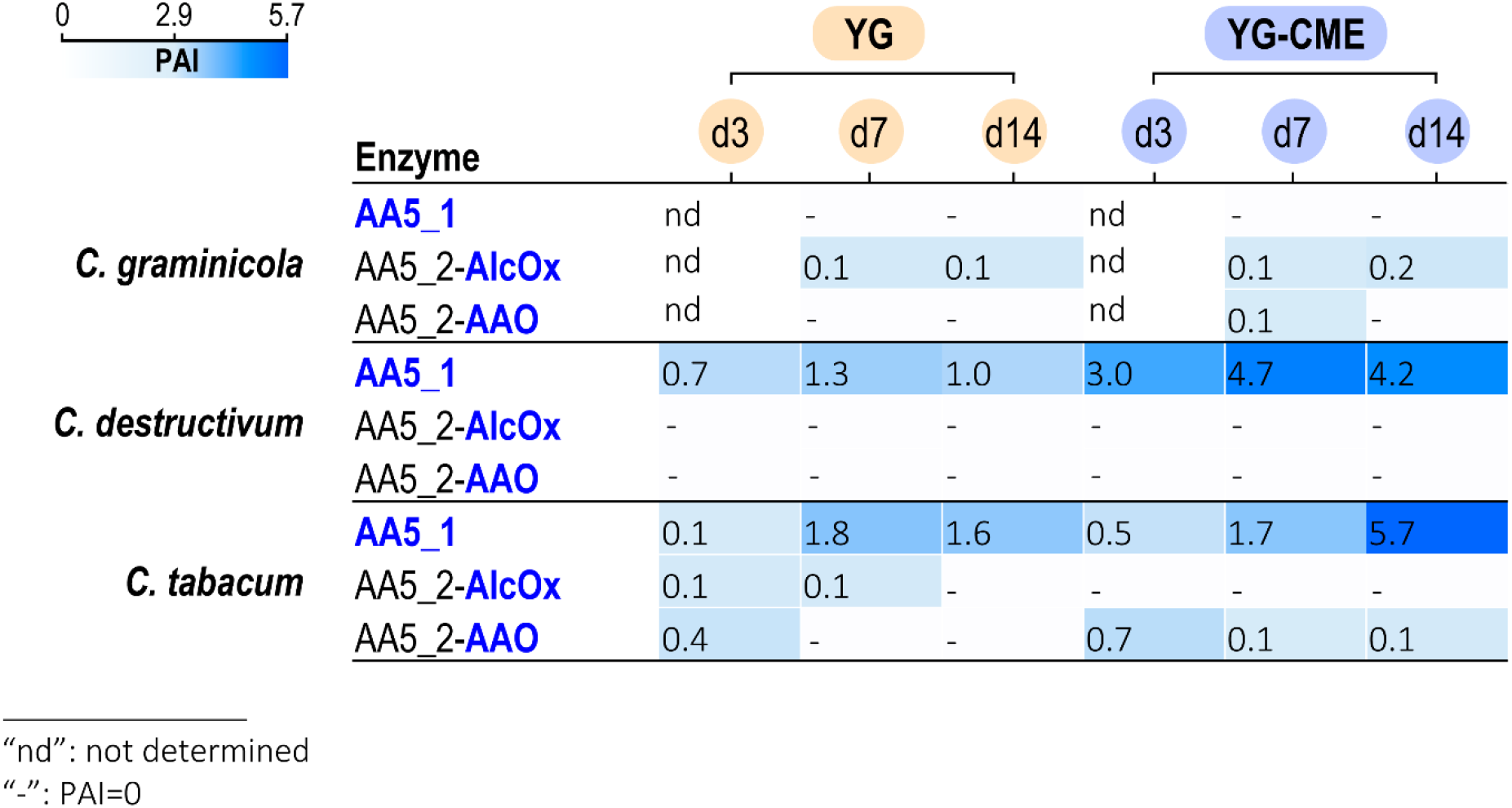
Heatmap of the abundance of AA5 detected in the fungal secretomes.

Together, these data provide unexpected information on the induction of secreted copper-enzymes in *Colletotrichum* species. However, the low level (or even the absence for *C. destructivum*) of AA5_2 detected under these experimental conditions indicates that the direct implementation of such secretomes for the bioconversion of fatty alcohols is not a viable strategy. These results prompted us to produce the *Colletotrichum* CRO-AlcOx enzymes recombinantly in *Pichia pastoris* and to test their ability to convert alcohols of interest for the flavor and fragrance industries. The objective was to find CRO-AlcOx with potential interesting traits not predictable by sequence analysis such as (i) activity on unactivated fatty alcohols, (ii) minimized requirement in accessory enzymes or (iii) reduced overoxidative activity.

### Recombinant production and substrate specificity of *Cde*AlcOx and *Cta*AlcOx

Detailed examination of the *Cde*AlcOx and *Cta*AlcOx sequences showed that the two AlcOx amino acid sequences share 90% identity with the *Cgr*AlcOx and likewise comprise a single catalytic module, as with all other characterized CRO-AlcOx from *Colletotrichum* spp. (Figure S1). The two AAO share 85% identity with the recently characterized *Cgr*AAO (44) and possess a N-terminal PAN_1 domain, in addition to the catalytic module, similarly to the *Cgr*AAO (Figure S1). We successfully produced both CRO-AlcOx in *P. pastoris* (Figure S6) with high titers for *Cta*AlcOx in flask culture (200 mg of purified enzyme per liter of culture) compared to *Cde*AlcOx (75 mg.L^-1^) and *Cgr*AlcOx (35 mg.L^-1^). Activity screening using the ABTS/HRP assay confirmed that both enzymes are indeed AlcOx and that they show similar substrate specificities as the canonical *Cgr*AlcOx (Figure 5). However, *Cde*AlcOx displayed a slightly higher specific activity on the secondary alcohol butan-2-ol, and on the longer aliphatic substrate decan-1-ol. This latter observation could be related to the M173L substitution compared to *Cgr*AlcOx, that could create a slightly more hydrophobic environment in the active site, favoring the accommodation of bulky alcohol substrates (Figure S7). The pH-rate profiles determined for *Cde*AlcOx and *Cta*AlcOx (Figure S8), revealed pH optima around 7.5 – 8 for both enzymes. For subsequent conversion assays and comparison with *Cgr*AlcOx, only *Cde*AlcOx was retained because *Cta*AlcOx exhibited no noticeable differences in terms of substrate specificity. Nevertheless, *Cta*AlcOx remains attractive from a biotechnological point of view, being 5.7-fold more highly expressed than the *Cgr*AlcOx in flask cultures.

**Figure 5:**
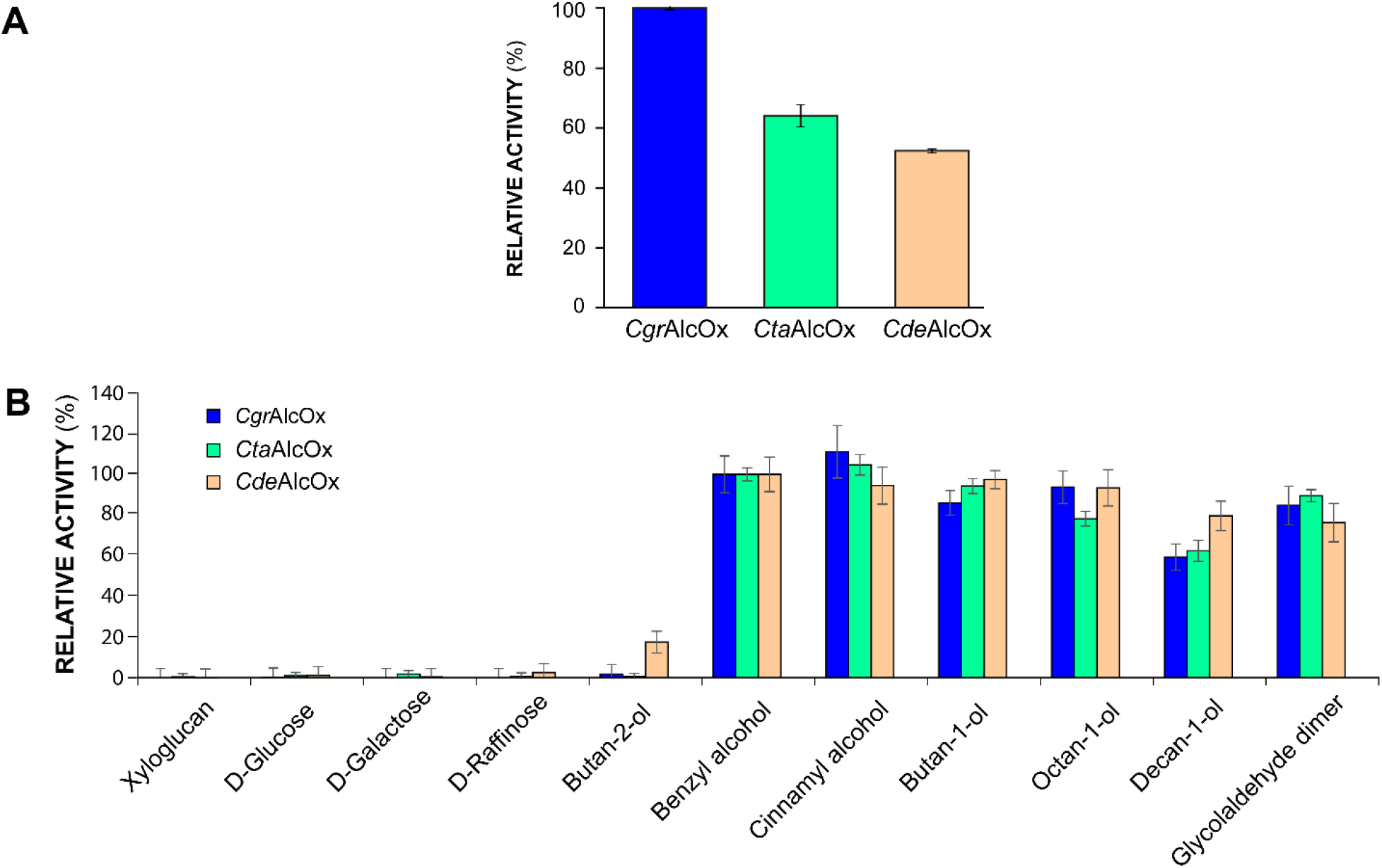
Comparative activity profiles of the three *Colletotrichum* AA5_2 AlcOx, **(A)** Relative activity of the recombinant AlcOx measured by ABTS/HRP coupled assay on 3 mM BnOH. **(B)** Relative activity profiling of the recombinant AlcOx enzymes on a set of substrates, determined by ABTS/HRP coupled assay, with BnOH set as reference (100%). Substrate concentrations were as follows: 1% (w/v) xyloglucan, 50 mM for D-glucose, D-galactose, D-raffinose, 3 mM for butan-2-ol, butan-1-ol, benzyl alcohol, cinnamyl alcohol, octan-1-ol, decan-1-ol, and glycolaldehyde dimer. Reactions were performed with 1 nM of *Cgr*AlcOx and *Cta*AlcOx, and 2 nM *Cde*AlcOx. Error bars show s.d. (independent experiments, n = 3).

### Biocatalytic production of odorant aldehydes using *Cde*AlcOx

*Cde*AlcOx was subsequently probed for the conversion of hexan-1-ol and octan-1-ol (two precursors of odorant aldehydes) using *Cgr*AlcOx as a benchmark. For these experiments, we tested the impact of the reaction time (15 minutes *versus* 16 hours conversion) and the addition of accessory enzymes (HRP and catalase). Conversion results showed that *Cde*AlcOx behave similarly to *Cgr*AlcOx with all the tested substrates (Figure 6). As previously observed with *Cgr*AlcOx (21, 33), *Cde*AlcOx alone was unable to promote full alcohol conversion. However, when both accessory enzymes were added to the reaction mixture, full conversion was readily achieved in 15 minutes. Carboxylic acid was not noticeable for hexan-1-ol but was observed after 16 hours of conversion of octan-1-ol only when HRP was present in the reaction. Indeed, this overoxidation to the acid was shown previously to be mediated by HRP in the case of fatty aldehydes > C6 (21). In the same study, we observed that in the case of benzaldehyde derivatives (i.e. 4-NO_2_-benzyl alcohol) some overoxidation was also detected but mediated by the *Cgr*AlcOx itself and dependent according to the propension of the aldehyde product to undergo hydration (21). Therefore, we included 4-NO_2_-benzyl alcohol in the panel of substrates (Figure S9) to probe whether *Cde*AlcOx behave similarly that *Cgr*AlcOx with aromatic *gem*-diols. The conversion of 4-NO_2_-benzyl alcohol indeed led to the corresponding acid after 16 hours of reaction even when HRP is omitted from the reaction. Altogether, these results confirm that the behavior observed for the *Cgr*AlcOx regarding accessory enzymes and overoxidation (21) applies to other CRO-AlcOx enzymes.

**Figure 6:**
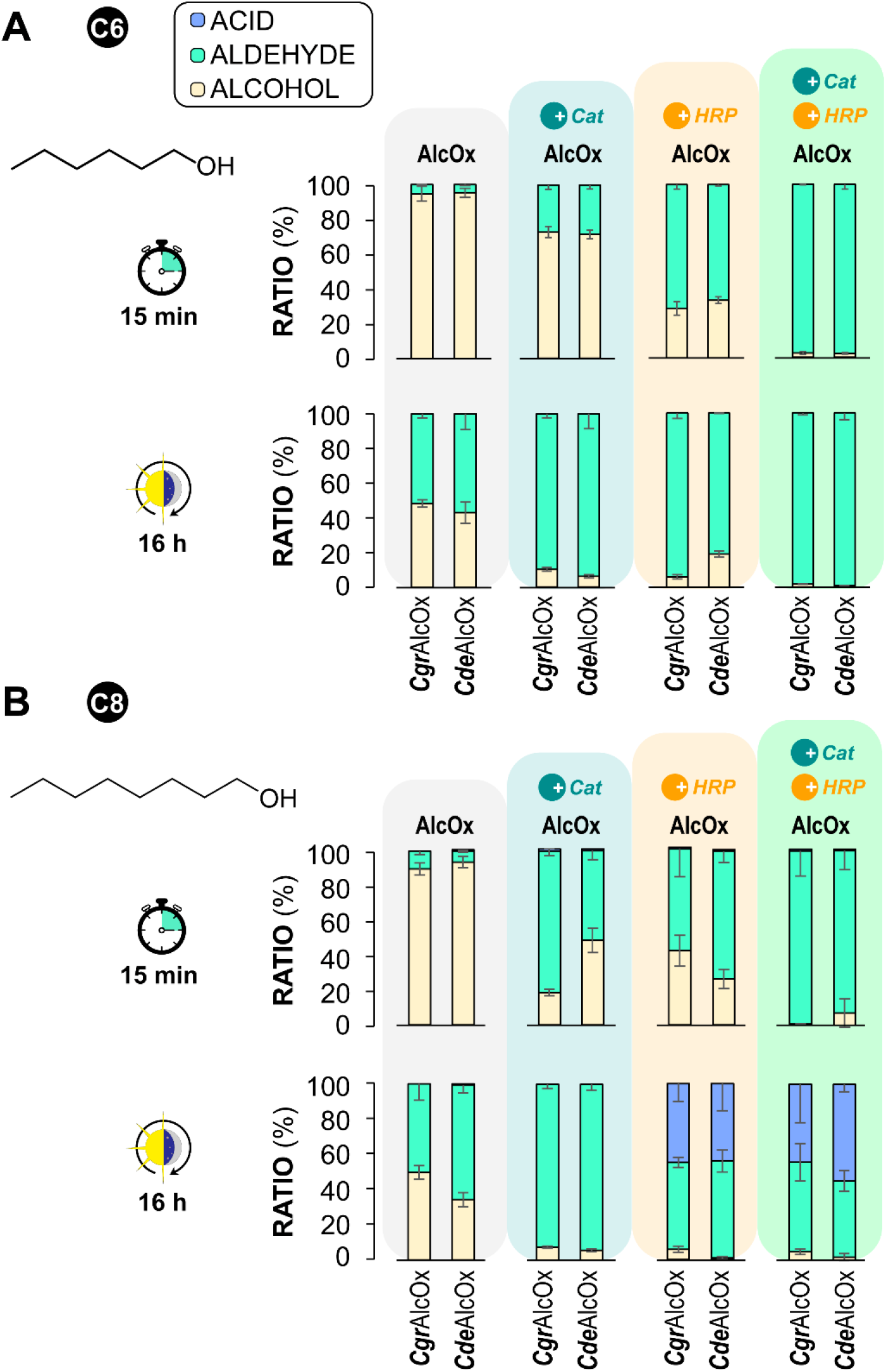
*Cgr*AlcOx and *Cde*AlcOx mediated oxidation of (A) hexan-1-ol and (B) octan-1-ol. Reactions were incubated for either 15 minutes or 16 hours. All reaction mixtures contained: 3 mM substrate and 1 µM AlcOx, in phosphate sodium buffer (50 mM, pH 8.0) and 23°C. Reactions varied as follows: no accessory enzyme added, addition of catalase (8 µM final), addition of HRP (12 µM final), addition of both catalase (0.5 µM) and HRP (12 µM final). Legend in panel A applies for panel B likewise. Conversion products were analyzed by GC-FID. Error bars show s.d. (independent experiments, n = 3).

## DISCUSSION

The time-resolved proteomic analyses performed here on *Colletotrichum* secretomes revealed interesting and unexpected results. We detected up to ca. 400 individual enzymes in the secretomes, with the maximum number found in the YG-CME medium. The high proportion of CAZymes in the secretomes is surprising considering the absence of any inducer to mimic plant cell walls. Strikingly, the addition of CME to the medium seemed to boost the secretion of AA enzymes. Moreover, our finding that some LPMOs (especially AA9) preferentially accumulated in the CME medium, in the absence of any plant biomass raises questions about the molecular basis of the regulatory pathways driving their production, and possibly their true biological function (52). In line with this observation, recent work suggests that some fungal LPMOs and LPMO-like proteins may be involved in biological functions distinct from plant biomass conversion, i.e., copper homeostasis and pathogenicity (53, 54) as well as fungal cell fusion (55). Other copper-containing AA enzymes exhibited higher accumulation in YG-CME cultures. Induction of the secretion of laccases could either be due to the addition of ethanol to the medium (12), or a response to oxidative stress (56). Indeed, it is known that at high concentrations, copper contributes to generate reactive oxygen species by Fenton chemistry (57, 58).

The proteins secreted exclusively in YG-CME medium revealed a range of chitin-active enzymes, as well as non-enzymatic chitin-binding proteins (LysM proteins) some of which are released by fungal cells upon interaction with their plant hosts, to conceal their presence from the host immune system by preventing fungal chitin recognition by the plant or protecting fungal hyphae against plant chitinases (59, 60). In the absence of the host or of any plant biomass, this observation suggests that the high concentration of CME in the medium could trigger some pathways related to pathogenesis. The presence of chitin active enzymes may also be indicative of a nutrient recycling in older senescing portions of the mycelium or of some fungal cell wall remodeling activity (61).

The relatively high abundance of GLOX from the AA5_1 subfamily in some YG-CME secretomes and the low secretion levels of AA5_2 AlcOx suggest distinct biological functions for these two types of CRO. AA5_1 GLOX are often described as enzymes acting on lignin in cooperation with some peroxidases in wood-decaying fungi (62). However, their high abundance here, under CME conditions, despite the absence of lignin points toward a possible additional biological function of GLOX in Ascomycota, related to filamentous growth or pathogenicity, as already demonstrated for the AA5_1 GLOX from the maize pathogen *Ustilago maydis* (63).

The primary objective of this study was to investigate the secretion of CRO-AlcOx by natural fungal strains with bioconversion potential. Under our experimental conditions, CRO-AlcOx are only sporadically secreted. Of note, a separate study very recently investigated the biological function of these enzymes and showed that they are co-secreted at low levels, over a short period of time during penetration of the plant host cuticle, and spatially restricted to the puncturing site (20). Thus, with hindsight, the secretion of large amount of CRO-AlcOx might be difficult to achieve without a proper signal to mimic the presence of plant host. Another option would be to identify and engineer the promotor region controlling co-expression of the genes encoding both the AlcOx and the associated peroxidase. However, for biocatalytic purposes, the recombinant expression of CRO-AlcOx remains the most promising alternative. We here confirmed that the knowledge gathered on sequence-structure-function of AA5_2 (18, 21, 44), can predict an AlcOx activity based on bioinformatic analysis and that various CRO-AlcOx can be easily produced heterologously using the yeast *P. pastoris* as host. *Cta*AlcOx was produced to high yield, which is promising for future upscaled production. The conversion assays performed with *Cde*AlcOx revealed that this enzyme can be used as a biocatalyst for the conversion of fatty alcohols. The behavior of *Cde*AlcOx in terms of overoxidation and accessory enzyme requirements suggests that the traits observed here and previously (20, 21, 33) are probably valid for all CRO-AlcOx.

In conclusion, the work carried out in this study highlights that *Colletotrichum* species can secrete a range of copper-containing oxidoreductases from the AA class without any plant cell wall inducer. While CRO-AlcOx were detected in some of the secretomes, the direct utilization of fungal strains for bioconversion of fatty alcohols will require further efforts to achieve a high secretion yield of CRO-AlcOx and a deeper understanding of the factors controlling their secretion. The recombinant pathway therefore represents the most promising approach to develop their use in biocatalytic applications for the production of fragrant fatty aldehydes.

## MATERIAL AND METHODS

### General information

All chemicals used for cultivation of fungi and microorganisms were reagent grade or higher. All chemicals used for biochemical assays were ≥ 95% purity. HRP type II and catalase from bovine liver were purchased from Sigma-Aldrich (Hamburg, Germany). Molar concentrations of HRP (MW: 33.89 kDa) and catalase (MW of monomer: 62.5 kDa) were estimated by Bradford assay using BSA as standard. Pre-cast 10% polyacrylamide gel Mini-PROTEAN TGX were purchased from Bio-Rad (USA).

### Data availability

The protein sequences of the *Cde*AlcOx and *Cta*AlcOx were deposited in GenBank under the accession numbers MZ269520 and MZ269521 respectively.

### Natural and recombinant strains

The following fungal strains were used: *C. graminicola* (M1.001, CBS 130836, (35)), *C. destructivum* (LARS 709, CBS 520.97, (64)) and *C. tabacum* (N150, CPC 18945, (64)). Fungi were maintained on Potato Dextrose Agar (PDA) plates, grown at 23°C for seven days and stored at 4°C before utilization. The recombinant *Pichia pastoris* X33 strains containing *Cgr*AlcOx gene (GenBank EFQ30446.1) was recovered from previous work (18, 21). DNA cloning and transformation of *P. pastoris* X33 with codon-optimized genes (for expression in *P. pastoris*) encoding *Cde*AlcOx and *Cta*AlcOx was carried out as described previously (65). The selection of the most productive recombinant transformants was done through culture and expression in 24-deepwell plates with subsequent purification from supernatant by affinity chromatography on Ni-NTA resin using an in-house automated procedure (65).

### Liquid culture and secretomes harvesting

Inocula were prepared by aseptically homogenizing the content of a 90 mm petri dish cultures of *Colletotrichum* (containing fungal biomass and PDA medium) using an Ultra Turrax T25 homogenizer (IKA, Germany) for 1 min at 9,500 rpm in 80 mL of the culture medium (see below). 200 μL of this homogenate was used to inoculate 100 mL of liquid medium placed in 500 mL baffled flask.

Liquid cultures were grown in two different media, namely: “YG” and “YG-CME”. YG medium was composed of glucose (10 g.L^-1^), ammonium tartrate (2 g.L^-1^), yeast extract (1 g.L^-1^), K_2_HPO_4_ (1 g.L^-1^), MgSO_4_ •7H_2_O (0.5 g.L^-1^), KCl (0.5 g.L^-1^) and 0.5 mL.L^-1^ of trace elements stock solution composed of (NH_4_)_6_Mo_7_O_24_•4H_2_O (20 mg.mL^-1^), FeSO_4_•7H_2_O (100 mg.mL^-1^), ZnSO_4_•7H_2_O (140 mg.mL^-1^), B_4_O_7_Na_2_•10H_2_O (200 mg.mL^-1^). Glucose was autoclaved separately as a 10X stock solution. The trace elements stock solution was sterilized by filtration through a 0.22 µm polyether sulfone membrane (Merck-Millipore, Germany). Note that insoluble elements present in this solution were removed by the filtration step. YG-CME medium was prepared exactly as YG but was further supplemented with CuSO_4_•5H_2_O (500 µM), MnCl_2_ (500 µM) and 9 g.L^-1^ ethanol, filtered sterilized (except for ethanol).

The inoculated cultures were placed in an Innova 42R incubator (New Brunswick, USA) and let grown at 25°C under natural light and shaking (at 150 rpm for *C. destructivum* and 105 rpm for *C. tabacum* and *C. graminicola*). The secretomes were harvested at day 3, day 7 and day 14 after inoculation. All samples were prepared in triplicates which were pooled upon harvesting. Each harvested secretome was submitted to several filtration steps: (i) Miracloth (Merck-Millipore), (ii) several glass microfiber filters used sequentially: 47 mm Whatman® grade GF/D, A and F (GE Healthcare, USA), (iii) 0.45 and 0.22-μm polyether sulfone membranes (Merck -Millipore) and finally diafiltered onto a polyether sulfone membrane with a 10kDa cut-off (Vivaspin, Sartorius, Göttingen, Germany) with 50 mM sodium phosphate buffer pH 7.0 to a final volume of 1-7 mL. The secretomes were stored at 4°C until use. Their total protein concentration was evaluated by the Bradford method (Protein Assay, BioRad, France) using a bovine serum albumin (BSA) standard protein.

### Microfluidic capillary gel electrophoresis

The concentrated secretomes samples were diluted so that 87.5 µL of the initially harvested secretomes (before concentration) were submitted to electrophoresis, except for *C. tabacum* YG-CME day 14 which was further diluted by 1/3 to avoid too high protein concentration. Samples were prepared in a 96-well plate using the manufacturer “HT Protein Express Assay Sample Buffer” (PerkinElmer, USA) supplemented with 36.4 mM (final concentration) of dithiothreitol. The samples were boiled at 95°C for 10 min and subjected directly to LabChip GX II capillary gel electrophoresis (PerkinElmer) according to the manufacturer’s instructions. The electropherograms were analyzed using the LabChip GX II software (PerkinElmer).

### LC–MS/MS protein identification

For each secretome, 15 μg of proteins were loaded on a 10% Tris–glycine precast SDS-PAGE gel (Mini-PRO-TEAN TGX, BioRad). After a short migration (0.5 cm) in the stacking gel, the gels were stained with Coomassie blue (BioRad) and each electrophoresis track was cut into two 2-mm-wide strips. Proteomic identification was performed at the Plateforme d’Analyse Protéomique de Paris Sud-Ouest (PAPPSO, INRA, Jouy-en-Josas, France; http://pappso.inra.fr/), according to the protocol described in (66). Briefly, the digestion of the proteins contained in the gel strips was carried out according to a standard trypsinolysis process, using modified trypsin (Promega, France). Peptide analysis was performed by a NanoLC Ultra 2D system (Eksigent Technologies, United States) coupled to a Q-exactive mass spectrometer (Thermo Fisher Scientific, France) using electro-spray ionization. Peptide attribution and protein annotation were performed by comparing mass spectrometry data to predicted proteins in the genomes of *Colletotrichum graminicola* M1.001 (https://mycocosm.jgi.doe.gov/Colgr1/Colgr1.home.html) and *Colletotrichum higginsianum* IMI 349063 (https://mycocosm.jgi.doe.gov/Colhig2/Colhig2.home.html) for both *C. destructivum* and *C. tabacum strains*. The internal contaminant database X!TandemPipeline software (X!TandemPipeline version 0.2.30, France) was also used. The protein annotation was completed manually using the CAZy database for the CAZymes annotation and using the BlastP tool from the NCBI database (https://blast.ncbi.nlm.nih.gov) for other proteins.

### Heterologous expression and purification

Recombinant *P. pastoris* X33 strains were streaked on yeast extract peptone dextrose (YPD) agar plates containing Zeocin (100 µg.mL^-1^) and incubated 3 days at 30°C. Five mL of YPD broth, in 50-mL sterile conical tubes, were inoculated with a *P. pastoris* transformant and incubated for five hours (30°C, 160 *rpm* in an orbital shaker). The preculture was used to inoculate 0.2% (*v/v*) of 500 mL buffered complex glycerol medium (BMGY) in a 2-liter flask, incubated (16 hours) until the OD600 nm reached 4-6. Then, the biomass was harvested by centrifugation (10 minutes, 16°C, 5,000 x *g*), resuspended in 100 mL of buffered complex methanol medium (BMMY) supplemented with CuSO_4_ (500 µM) and methanol (1% *v/v*) in a 500 mL flask, and shaken for 3 days in an orbital shaker (200 *rpm*, 16°C) with daily feeding of methanol (1% *v/v*). The culture was then centrifuged (10 minutes, 4°C, 5,000 x *g*) and the supernatant containing the secreted proteins, filtrated on a 0.45-µm cut-off membrane (Millipore, USA) followed by buffer exchange with Tris-HCl (50 mM, pH 8.5), performed by ultrafiltration through a 10 kDa cut-off polyether sulfone membrane (Vivacell 250, Sartorius Stedim Biotech GmbH, Germany). The buffered supernatant was loaded on a DEAE-20 mL HiPrep FF 16/10 anion exchange chromatography column (GE Healthcare), equilibrated with Tris-HCl buffer (50 mM, pH 8.5) and connected to an Äkta Xpress system (GE Healthcare). Elution was performed by applying a linear gradient from 0 to 500 mM NaCl (in Tris-HCl buffer 50 mM, pH 8.5) with a flow rate set to 5 mL.min^-1^. The collected fractions were analyzed by SDS-PAGE (10% polyacrylamide precast gel, Bio-Rad) stained with Coomassie blue. Fractions containing the recombinant enzyme were pooled, concentrated and exchanged to sodium phosphate buffer (50 mM, pH 7.0). The *Cgr*AlcOx (ε^280^ = 101,215 M^-1^.cm^-1^), *Cde*AlcOx (ε^280^ = 94,225 M^-1^.cm^-1^) and *Cta*AlcOx (ε^280^ = 94,225 M^-1^.cm^-1^) concentration was determined by UV absorption at 280 nm using a Nanodrop ND-200 spectrophotometer (Thermo Fisher Scientific). Final solutions (3-5.5 mg.mL^-1^) were flash-frozen using liquid nitrogen and stored at -80°C for long-term storage or stored at 4°C for immediate use.

### Spectrophotometric enzyme activity assays

AlcOx initial rates were determined by monitoring H_2_O_2_ released during AlcOx oxidation of various substrates, using the coupled ABTS/HRP assay. Routine assays were performed in 96-wells transparent microtiter plates (Flat bottom, Polystyrene - Greiner Bio One, Austria) in 200 µL final volume containing 0,25 mg/mL ABTS powder, 0,1 mg.mL^-1^ HRP powder (as provided by the supplier), 3 mM substrate (unless indicated otherwise) and 1 nM AlcOx in sodium phosphate buffer (50 mM, pH 8.0), at 23°C. Reactions were initiated by the addition of substrate to a premix containing all other reagents. Evolution of the absorbance at 414 nm (ABTS cation radical) was measured over time with a Tecan Infinite M200 (Tecan, Switzerland) plate reader. Oxidation of 1 mole of substrate by AlcOx consumes 1 equivalent of O_2_ and generates 1 equivalent of H_2_O_2_ which is in turn used by HRP as co-substrate to oxidize 2 equivalents of ABTS. Insoluble substrates were prepared in acetone so that the final acetone concentration in the enzyme assay did not exceed 1% (*v/v)*. Standard curve of know concentrations of H_2_O_2_ (1 to 40 µM) were made and used to quantify H_2_O_2_ production. One unit of AlcOx activity was defined as the amount of enzyme necessary to produce 1 µmole of H_2_O_2_ per minute.

### Bioconversion assays

Bioconversions were carried out in 4 mL-clear borosilicate glass vials closed by screw caps with PTFE septum for a total reaction volume of 500 µL. Routine assay contained 3 mM substrate (prediluted in acetone), 1 µM AlcOx, various amounts of catalase (0-8 µM) and/or HRP (0-12 µM), in sodium phosphate buffer (50 mM, pH 8.0). Final reaction contained 1% (*v/v*) acetone. Reactions were run at 23°C, under stirring at 190 *rpm* in an Innova 42R incubator (New Brunswick, USA). Vials were placed lying down in the incubator. The reaction mixture was then acidified by addition of 10 µL HCl (12 M) and products and possible remaining substrate were extracted by adding 500 µL of cyclohexane/ethyl acetate mixture (1:1), followed by shaking and centrifugation for 5 minutes at 1.500 x *g*. The organic layer was transferred into a new vial by pipetting and analyzed with a GC-2014 apparatus (Shimadzu, Japan) equipped with a flame ionization detector (FID) and an Optima-σ-3 GC capillary column (30 m x 0.25 mm x 0.25 µm - Macherey-Nagel GmbH & Co, Germany). Nitrogen (200 kPa) was used as carrier gas. The injector and detector temperatures were set at 250°C and temperature programs are described in Table S2. Heptanal or guaiacol were used as internal standards for either aliphatic or aromatic products analyses, respectively.

## AUTHOR CONTRIBUTIONS

D.R. carried out most of the experimental work, analyzed the data. S.S. was involved in the culture of fungi and production of the secretomes. L.O.C. and D.N. performed proteomics and analysed data. E.D. and B.H. performed expert annotation of CAZymes. M.H. contributed to enzyme production in *Pichia pastoris*. S.G. performed the microfluidic capillary gel electrophoresis experiments. R.J.O. and N.L. provided the *Colletotrichum* strains and access to the associated genomic data. M.L. and J-G.B conceptualized the study. F.L., B.B., M.L. and J-G.B. supervised the work. D.R. and J-G.B. drafted the manuscript. All authors approved the final version of the manuscript.

## ACKNOWLEDGEMENTS

This study was supported by the French National Agency for Research (“Agence Nationale de la Recherche”) through the “Projet de Recherche Collaboratif International” ANR-NSERC (FUNTASTIC project, ANR-17-CE07-0047). We are grateful to MANE & Fils and the “Association Nationale Recherche Technologie” (ANRT) for funding the PhD fellowship of D.R. entitled “Discovery and structure-function study of new fungal copper radicals oxidases used as biocatalysts for the valorisation of alcohols and plant biomass”. This *Convention Industrielle de Formation par la RecherchE* (CIFRE) grant no. 2017/1169 runs from 1 April 2018 to 1 April 2021. Fungal genome sequencing was supported by funding to R.J.O. (grant ANR-17-CAPS-0004-01).

We are grateful to Renaud Vincentelli (AFMB – CNRS – Marseille, France) for providing access to the microfluidic capillary gel electrophoresis apparatus.

